# Auxin-Mediated Sterility Induction System for Longevity and Mating Studies in *Caenorhabditis elegans*

**DOI:** 10.1101/284232

**Authors:** Katja R. Kasimatis, Megan J. Moerdyk-Schauecker, Patrick C. Phillips

## Abstract

The ability to control both the means and timing of sexual reproduction provides a powerful tool to understand not only fertilization, but also life history trade-offs resulting from sexual reproduction. However, precisely controlling fertilization has proved a major challenge across model systems. An ideal sterility induction system should be external, non-toxic, and reversible. Using the auxin-inducible degradation system targeting the *spe-44* gene within the nematode *Caenorhabditis elegans*, we designed a means of externally inducing spermatogenesis arrest. We show that exposure to auxin during larval development induces both hermaphrodite self-sterility and male sterility. Moreover, male sterility can be reversed upon cessation of auxin exposure. The sterility induction system developed here has multiple applications in the fields of spermatogenesis and mating systems evolution. Importantly, this system is also a highly applicable tool for aging studies. We show that auxin-induced self-sterility is comparable to the commonly used chemically-induced FUdR sterility, while offering multiple benefits, including being less labor intensive, being non-toxic, and avoiding compound interactions with other experimental treatments.

## INTRODUCTION

Sexual reproduction is among the most fundamental of biological processes, and the ability to control the means and mode of sexual reproduction provides a powerful tool for studying a wide variety of important questions. First and foremost, interactions within and between the sexes are mediated by the process of fertilization. Thus, precise control of fertilization allows for the nature of those interactions to be directly manipulated, addressing questions regarding potentially antagonistic interactions between members of the same sex (e.g., sperm competition) (Karr and Pitnick 1999; Edward *et al.* 2015), between members of the opposite sex (e.g., sexual conflict) (Arnqvist and Rowe 2005), and between parents and offspring (e.g., parent-offspring conflict) (Trivers 1972). In addition, reproduction itself is a critical field of study. For example, investment in offspring is often thought to represent a trade-off with other aspects of an individual’s life history, including overall lifespan (Stearns 1989; Schluter *et al.* 1991). Directly manipulating the dynamics of reproduction allows these trade-offs to be specifically assessed. More prosaically, some experiments, such as longevity studies, require the separation of parents and offspring and in some systems this separation is best accomplished by simply not allowing the adults to reproduce in the first place (Park *et al.* 2017).

Currently there are few techniques available to control reproduction short of direct physical manipulation of sexes. In these cases, mostly in model organisms, chemical interventions or genetic mutations can be used to induce sterility in one of the sexes. For example, sterility induction mechanisms—both genetic and chemical—are common in the agricultural industry as a method of preventing cross-pollination (Kempe and Gils 2011). In *Drosophila*, several genetic mutations can be used to generate either female (Schüpbach and Wieschaus 1991; Volpe *et al.* 2001) or male sterility (Castrillon *et al.* 1993). However, since these mutations tend to be recessive, they must be maintained over a balancer chromosome or in a heterozygous population, making them manually intensive to use. Vertebrate models offer many more challenges to reproductive control, and therefore few sterility induction approaches exist in these systems (see Hsu *et al.* 2009). *Caenorhabditis elegans* is a major model system for genetics, development, neurobiology, and aging. Within *C. elegans*, a limited number of sterility mutants are available (L’Hernault 2006; Ellis and Stanfield 2014) and can be maintained by mating hermaphrodites to males. In some cases, temperature sensitive sterility mutants (Hirsh and Vanderslice 1976; Ward and Miwa 1978) exist. However, these tend to be general germline mutations with pleiotropic effects rather than fine-scale control of gametogenesis *per se*. Additionally, these mutants by necessity require a temperature shift, which can affect lifespan (Park *et al.* 2017). An alternative scheme is to prevent progeny production in hermaphrodites using chemical treatments (Mitchell *et al.* 1979), though these techniques are manually intensive and not conducive to high-throughput assays. Further, chemical intervention can potentially generate unaccounted for fitness effects, which not only confound the biological interpretation of results but also make reproducibility a challenge.

An ideal sterility system would be inducible, driven by an external treatment, and, when possible, reversible. To the best of our knowledge such an approach does not exist, even within model organisms. To address this need, we used the non-toxic, non-native auxin inducible degradation (AID) system (Nishimura *et al.* 2009) coupled with knowledge of a critical spermatogenesis gene to create an external sterility induction system in *C. elegans*. We show that this system induces self-sterility of hermaphrodites and complete, but reversible sterility of males. This method has broad applications in nematode biology, including studies of aging, gametogenesis, and mating systems evolution.

### Constructing an inducible spermatogenesis arrest

The AID system (Nishimura *et al.* 2009; Zhang *et al.* 2015) was chosen as the optimal method for an external sterility induction system in *C. elegans*, as auxin is non-native, non-toxic, and cost-effective. The auxin hormone regulates gene expression in *Arabidopsis thaliana* by activating the F-Box transport inhibitor response 1 (TIR1) protein – the substrate recognition component of a Skp1-Cullin-F-box E3 ubiquitin ligase complex which ubiquitinates degron-tagged proteins for degradation by the proteasome (Tan *et al.* 2007; Nishimura *et al.* 2009). This system has been co-opted as an inducible genetic mechanism in a variety of organisms by degron-tagging a protein of interest and choosing a promoter to drive TIR1 expression in the necessary cell type (Kanke *et al.* 2011; Zhang *et al.* 2015; Trost *et al.* 2016; Natsume *et al.* 2016). We targeted a necessary spermatogenesis gene *spe-44*, causing a spermatogenesis arrest and therefore sterility. Specifically, *spe-44* is one of eleven sperm-specific transcription factors (Reinke 2003) and is predicted to have hundreds of downstream targets, including the critical Major Sperm Protein (Kulkarni *et al.* 2012). Constitutive TIR1 expression was driven using the germline promoter of *pie-1*, which is one of few genes known to have strong sperm expression in hermaphrodites and males (Merritt *et al.* 2008). These three components—auxin, *P*_*pie-1*_*::TIR1, spe-44::degron*—generate a fully controllable sterility induction system in *C. elegans.*

## MATERIALS AND METHODS

### Molecular biology

Guide sequences were chosen using the tools CRISPRdirect (Naito *et al.* 2015), MIT CRISPR Design (http://crispr.mit.edu) and Sequence Scan for CRISPR (Xu *et al.* 2015). For the TIR1 insertion, a guide targeting the sequence GAAATCGCCGACTTGCGAGG<u>AGG</u> near the ttTi4348 MosSCI site was inserted into pDD162 (Dickinson *et al.* 2013) using the Q5 site-directed mutagenesis kit (NEB) to create pMS18. This insertion region was previously shown to be permissive for germline expression (Frøkjær-Jensen *et al.* 2012). The plasmid pMS30 was created by Gibson assembly using the NEBuilder HiFI Kit (NEB) and included: homology arms amplified from N2 genomic DNA, the *pie-1* promoter amplified from pCM1.127 (Addgene #21384) (Merritt *et al.* 2008), the *C. elegans* optimized *AtTIR1::mRuby* fusion and *unc-54* terminator amplified from pLZ31 (Addgene #71720) (Zhang *et al.* 2015), and the self-excising drug selection cassette (SEC) amplified from pDD282 (Addgene #66823) (Dickinson *et al.* 2013). The plasmid backbone was also derived from pDD282. An 11 bp segment of the genomic DNA sequence was omitted from the homology arms to prevent re-cutting. All plasmid assembly junctions were confirmed by Sanger sequencing. Sequencing showed that pMS30 contained a single nucleotide substitution in one of the LoxP sites of the SEC. However, this substitution did not significantly impact SEC removal.

The degron:3X-FLAG tag utilized asymmetric homology arms (Richardson *et al.* 2016) for *spe-44* insertion and contained appropriate silent sites to prevent re-cutting. The insert was synthesized as a GeneArt String (ThermoFisher) and amplified by PCR prior to injection.

### Strain generation by CRISPR/Cas9

The *P*_*pie-1*_*::TIR1::mRuby* construct was injecting into the gonad of young adult hermaphrodites (standard laboratory strain N2) using a mixture of 50 ng/μl pMS18, 10 ng/μl pMS30 and 2.5 ng/μl pCFJ421 (Addgene #34876) (Frøkjær-Jensen *et al.* 2012). Screening and removal of the SEC was done following Dickinson *et al.* (2013). Presence of the insertion and removal of the SEC was confirmed by PCR and Sanger sequencing.

To degron tag *spe-44*, a cr:tracrRNA (Synthego) targeting the sequence ATTGAATATGACTAGGTCC<u>TGG</u> near the C-terminus of *spe-44* was annealed and pre-incubated with Cas9 (PNA Bio) in accordance with manufacturer protocol. A mix of 1.7 μM cr:tracrRNA, 1.65 μg/μl Cas9, and 80 ng/μl of the PCR repair template, was then injected into the gonad of young adult N2 hermaphrodites containing the *P*_*pie-1*_*::TIR1::mRuby* construct. Included in the injection mix was an additional cr:tracrRNA and oligonucleotide repair template, allowing for screening through *dpy-10* co-conversion (Paix *et al.* 2015). Progeny from broods containing individuals with a dumpy or roller phenotype were then screened for the *spe-44::degron* insertion by PCR and confirmed by Sanger sequencing.

Confirmed double mutants were backcrossed 5 times to N2 to create the final strain PX627 (fx*Is*1[*P*_*pie-1*_*::TIR1::mRuby*, I:2851009]; *spe-44*(fx110[*spe-44*::*degron*]). This strain was crossed to strain CB4088, to create the male-rich strain PX629 (fx*Is*1[*P*_*pie-1*_*::TIR1::mRuby*, I:2851009]; *spe-44*(fx110[*spe-44*::*degron*]) IV; *him-5* e(1490) V).

### Worm culture and strains

The *C. elegans* strains PX627, PX629, N2, and JK574 (*fog-2*(q71) V) were maintained on NGM-agar plates seeded with OP50 *Escherichia coli* at 20°C (Brenner 1974). The *fog-2* mutation blocks self-sperm production in hermaphrodites, making them a functionally female. Synchronized cultures of larval stage 1 (L1) animals were obtained through hypochloride treatment of gravid adults (Kenyon 1988). To induce sterility, worms were transferred to NGM-agar plates containing 1 mM indole-3-acetic acid (Auxin, Alfa Aesar) following Zhang *et al.* (2015). Zhang *et al.* (2015) showed this auxin concentration to be non-toxic to adults with no larval development defects or fecundity effects. Auxin plates were stored in the dark at 4°C to prevent compound degradation.

All larval exposure assays were carried out on small plates (35 mm) seeded with 100 μL *E. coli* and a sample size of 130 hermaphrodites per developmental stage and 100 males per stage. Adult male developmental exposure assays were done by plating synchronized L1 PX629 worms on NGM-agar plates until day 1 of adulthood. Males were then transferred to small auxin plates seeded with 10 μL *E. coli* along with two virgin females (strain JK547). Males were transferred to new virgin females twice a day until no fertilized eggs were seen on plates. Male sterility recovery experiments were done by plating synchronized L1 animals on auxin plates and leaving worms on auxin until day 1 or day 2 of adulthood. Males were then transferred to small NGM plates seeded with 10 μL *E. coli* and given three virgin females (strain JK574) with which to mate. Plates were monitored until fertile eggs appeared.

For the hermaphrodite self-sterility mating experiments, synchronized L1 PX627 worms were plated onto auxin plates and removed three hours into adulthood. Virgin females (strain JK574) were used as a control. Individual pseudo-females were mated with two males (strain JK574) overnight on small NGM-agar plates seeded with 10 μL *E. coli*, after which males were removed. All the progeny laid over the subsequent 24 hours were counted. Two independt biological replicates were done with 23 to 35 pseudo-females in each treatment. Fertility data were analyzed using a generalized linear model (GLM) framework with random effects and a Poisson distribution using the *lme4* v.1.13 package (Bates *et al.* 2015) in the R statistical language (R Core Team 2015).

### Lifespan assays

Lifespan data were collected using automated lifespan machines following Stroustrup *et al.* (2013). Briefly, worms were synchronized by letting day 2 adults (strains PX627 and N2) lay eggs over a two hour time period. Auxin self-sterility was achieved by allowing PX627 hermaphrodites to lay directly on auxin plates or by transferring larval stage 4 (L4) progeny to auxin plates. Both self-sterility treatments were transferred to NGM-agar plates on day 1 of adulthood. As a control, egg lays for both PX627 and N2 hermaphrodites were done on NGM-agar plates. At day 1 of adulthood, these animals were transferred to small plates containing 51 μM 5-fluoro-2’-deoxyuridine (FUdR, VCI America) to inhibit reproduction (Mitchell *et al.* 1979). Control worms were transferred to fresh FUdR plates 24 hours later.

On day 5 of adulthood, all worms were transferred onto medium scanner plates (60 mm) with sealable lids to minimize dehydration. NGM-agar scanner plates contained 40 mM potassium phosphate buffer (pH 6.0), 1mM magnesium sulfate, and 5 mg/mL cholesterol, along with 100 mg/mL nystatin to prevent fungal growth while on the automated lifespan system. Control plates also included 51 μM FUdR. All scanner plates were seeded with 200 μL *E. coli*. A total of 35 to 60 adult hermaphrodites were transferred to each plate with four technical replicates of each treatment. The 16 plates were randomly arranged on a modified Epson v700 scanner in a temperature controlled 20°C room and held in place by rubber mat. Plates were imaged approximately every hour for twenty days across two independent biological replicates.

Images were analyzed using the Worm Browser software developed with the automated lifespan system (Stroustrup *et al.* 2013). This process includes specifying the location of individual plates on the scanner, detecting individual worms, and analyzing worm movement. The resulting data are time of death calls for each individual worm based on the cessation of movement. All plates were hand annotated to ensure that non-worm objects were excluded. Additionally, the time of death calls for the first and last 10% of worms on each plate were checked as these time points are more error prone. The final lifespans were calculated using the egg lay as day zero.

To analyze the influence of our sterilization approach on longevity, we used a mix-model survival analysis as outlined in Lucanic *et al.* (2017). Longevity effects were evaluated using both a mixed-model Cox Proportional Hazard (CPH) model (Therneau *et al.* 2012) using the *coxme* v.2.2-5 package {coxme:DjUq3nzA}, as well as via GLM using the *lme4* package in R. In each case, the coxme and GLM approaches yield equivalent results and so only the *coxme* results are presented as they represent the more comprehensive analytical framework for these data. Using the automated lifespan machine, a small subset of individuals initially placed on a plate are missing and presumed lost over the course of an assay. Such individuals would normally be classified as “censored” in normal survivorship analysis. However, because mortality is determined retrospectively when an individual ceases to move, the moment of loss of such individuals cannot be determined and so they must simply be classified as missing rather than censored at a given time point. For these analyses, the environmental treatment within which each individual was raised (FUdR, auxin) and the genotype of the individual (wildtype or *P*_*pie-1*_*::TIR1::mRuby; spe-44::degron*) were treated as fixed effects, while replicate and plate (nested within replicate) were treated as random effects. Specific *a priori* hypotheses about effects of FUdR and genetic background were tested via contrast coefficients using the *mcp* procedure of the multcomp procedure in R (Hothorn *et al.* 2008).

### Data accessibility

The oligonucleotides and synthetic constructs used in this study are listed in Table S1. The fertility data are given in File S1 and the lifespan data in File S2. Worm strains N2, JK574, PX627, and PX629 are available from the *Caenorhabditis* Genetics Center.

## RESULTS

### Self-sterility induction in hermaphrodites

*Caenorhabditis elegans* hermaphrodites are protandrous, such that they produce several hundred sperm cells during their final larval stage and then switch to oocyte production for the remainder of their lifespan (Hirsh *et al.* 1976). We examined the necessary and sufficient windows of auxin exposure during hermaphrodite development to induce self-sterility. To prevent sperm production, hermaphrodites must be exposed to auxin during their larval development (Fig. 1A). Specifically, the L4 window alone—the developmental stage during which sperm are produced—was both necessary and sufficient to drive self-sterility (Fig. 1B). As observed in other spermatogenesis mutants, self-sterile hermaphrodites continue to lay unfertilized oocytes throughout their adult life. Adult exposure to auxin had no effect on progeny production.

**Figure 1.**
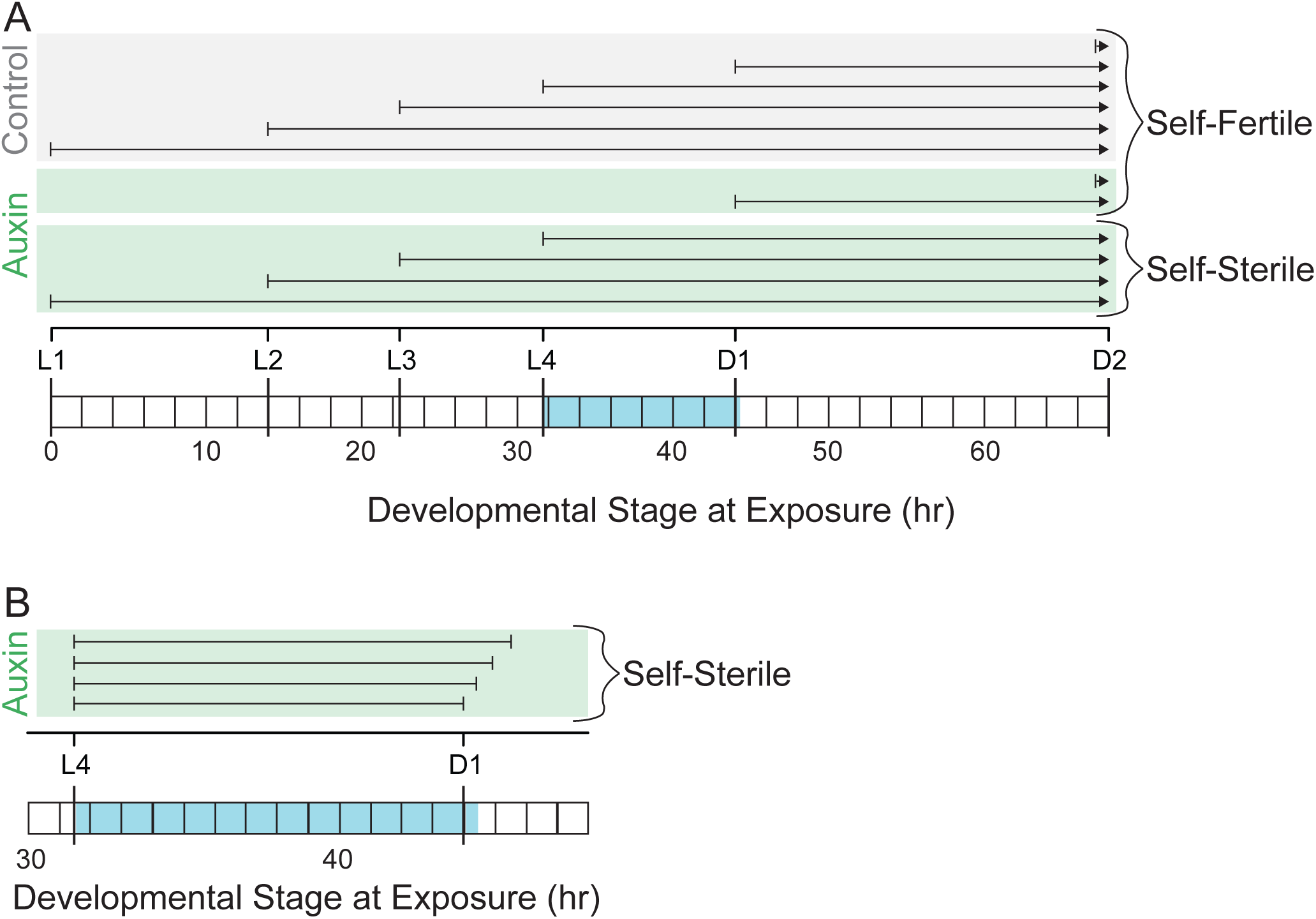
Auxin exposure induces hermaphrodite self-sterility. A) Hermaphrodites were exposed to auxin (shown in green; n = 130 hermaphrodites per stage) starting at each of the four larval stages (L1 – L4) and first two days of adulthood (D1 – D2). Spermatogenesis (highlighted in blue) occurs during L4 and continues approximately 30 minutes into adulthood (Hirsh *et al.* 1976). When exposed to auxin during larval development, all hermaphrodites were self-sterile. However, adults exposed to auxin were fully fertile. B) Hermaphrodites were exposed to auxin at L4 and transferred off auxin in 30 minute increments starting at adulthood (n = 50 hermaphrodites per stage). All worms were self-sterile, indicating that L4 alone is sufficient to induce self-sterility.

Despite being self-sterile, hermaphrodites could recover their fertility when mated to a male, comparable to hermaphrodites made functionally female through the *fog-2* mutation (Fig. 2). Interestingly, these mated self-sterile hermaphrodites were highly consistent in the number of progeny they produced. However, control *fog-2* females laid significantly more progeny than self-sterile hermaphrodites (z = −2.78, p < 0.01), likely due to adaptation to obligate outcrossing.

**Figure 2.**
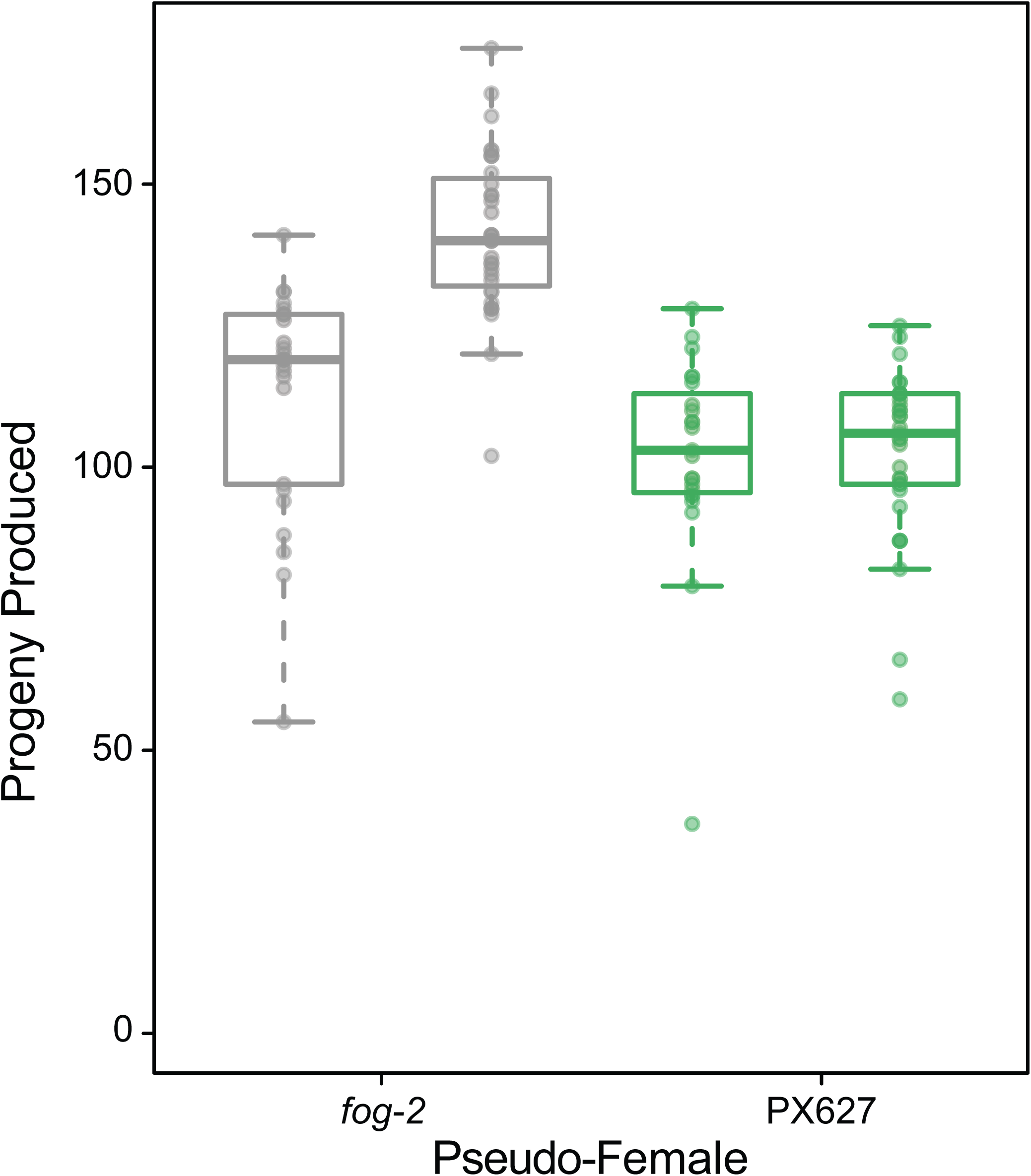
Self-sterile PX627 hermaphrodites (green) can recover their fertility when mated with a wildtype male, as compared to *fog-2* functional females (gray). Each bar within a genotypic set represents an independent replicate. While self-sterile hermaphrodites produced fewer progeny than *fog-2* females (z = −2.78, p < 0.01), their progeny production was invariable across replicates (t = −0.097, df = 40.73, p = 0.92).

### Inducible sterility of males is reversible within a single generation

We tested the sterility induction of males using a male-enriched *C. elegans* strain. Like hermaphrodites, males begin spermatogenesis during L4, however they continue producing sperm throughout adulthood, whereas hermaphrodites do not (L’Hernault 2006). We examined the window of auxin exposure during larval male development sufficient to induce sterility. Interestingly, L4 exposure alone was not sufficient to induce complete sterility, as these males still produced a low number of progeny. Rather males had to be exposed to auxin at least 2 hours prior to the L3/L4 molt (Fig. 3A).

**Figure 3.**
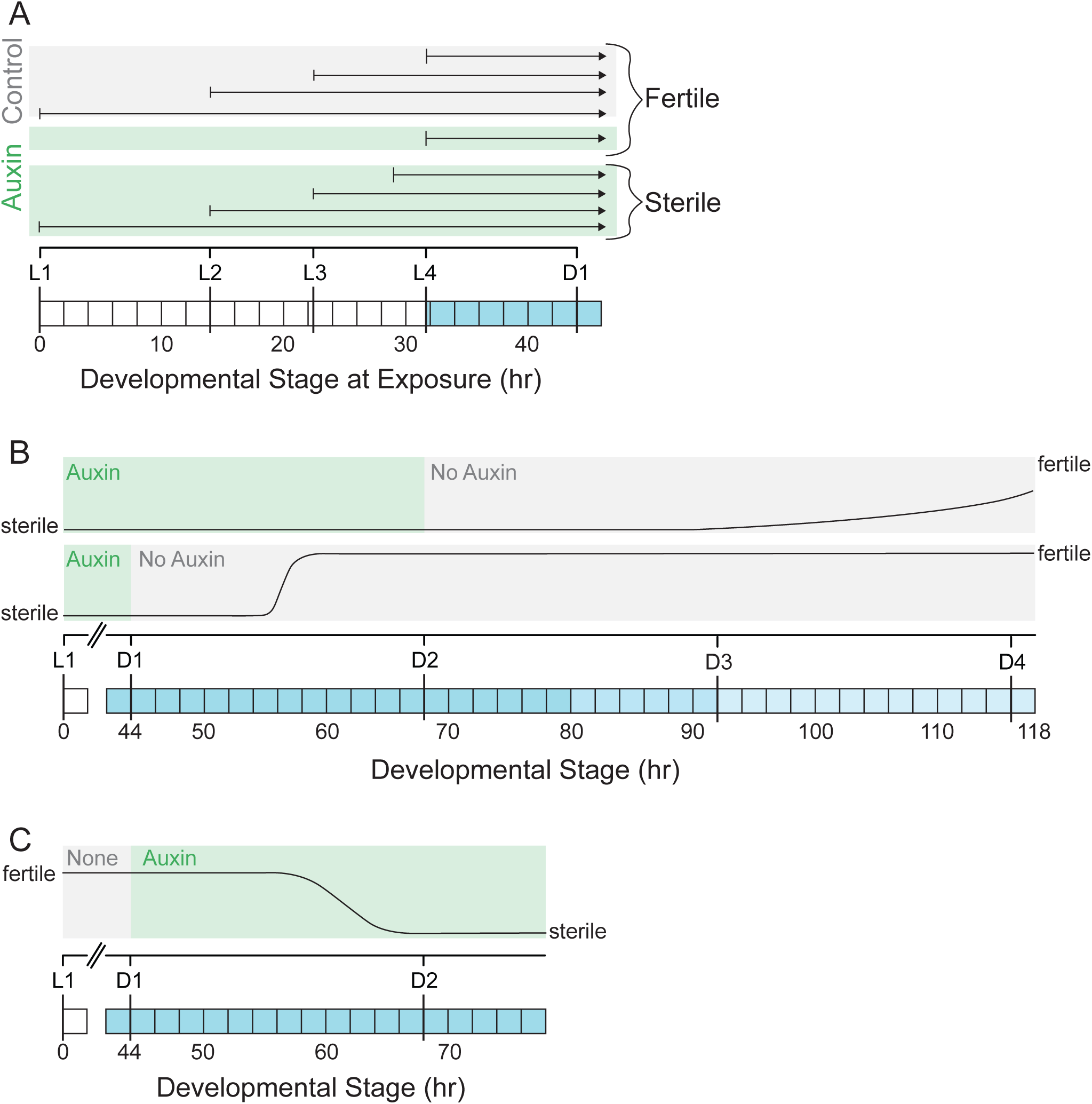
Auxin exposure induces male sterility. A) Males were exposed to auxin (shown in green; n = 100 males per stage) during larval development. Spermatogenesis (highlighted in blue) begins during L4 and continues throughout adulthood (L’Hernault 2006). Males exposed to auxin prior to L4 were sterile, however, males exposed to auxin at L4 produced a low number of progeny. B) Males exposed to auxin throughout larval development up until day 1 (D1; n = 30) or day 2 (D2; n = 30) of adulthood recovered their fertility when removed from auxin in a time-dependent manner. However, not all males removed from auxin at day 2 of adulthood became fertile (n = 16). C) Adult male sterility could be induced by exposing males to auxin at day 1 of adulthood. All males were sterile within 24 hours (n = 44).

To determine if sterility in males could be reversed following consistent exposure to auxin during larval development, males were transferred from auxin to standard NGM plates at day 1 and day 2 of adulthood. Day 1 adult males began to recover their fertility within approximately 12 hours and all males were fully fertile within 24 hours (Fig. 3B). Day 2 adult males, however, had a much slower recovery period and not all males became fertile (n = 16 out of 30). Alternatively, we measured the sterility induction onset for adult male auxin exposure. Here males were raised on standard NGM plates and exposed to auxin starting at day 1 of adulthood. Within 24 hours of auxin exposure, no progeny were observed from male-virgin female matings, indicating that males were fully sterile (Fig. 3C).

### Hermaphrodite self-sterility induction as a tool for aging research

Lifespan assays in *C. elegans* are complicated by the difficult and relatively labor intensive process of separating individuals of an aging cohort from their offspring. A variety of approaches to address this problem are used in the literature, with treatment of adults by the pyrimidine analog FUdR (a chemotherapy agent) being the most widely used. FUdR interferes with DNA synthesis, thereby preventing the production of viable offspring. The sterility induction system developed here allows hermaphrodites to be treated during larval development in order to induce self-sterilization and to then be transferred to whatever media type is necessitated by a given experiment, such as plates treated with bioactive compounds (see Lucanic *et al.* 2017). We examined this potential use by contrasting longevities in wildtype (N2) and *Ppie-1::TIR1::mRuby; spe-44::degron* (PX627) adults living on plates containing FUdR with those of PX627 individuals reared on auxin plates either throughout the entire larval development period or during the L4 stage alone before being transferred to standard NGM plates for the remainder of their lives.

Median lifespan of individuals sterilized via either FUdR or auxin were similar to one another, although they differ slightly in quantitative details (Fig. 4). A comparison of adult wildtype and PX627 individuals raised on FUdR in the absence of auxin yielded highly similar survivorship profiles and median lifespans (N2_FUdR = 18.1 days, PX627_FUdR = 17.5 days; CPH contrast: *z* = 2.17, *p* = 0.0996). PX627 individuals sterilized with FUdR also displayed quite similar overall longevity profiles, with individuals raised on auxin for their entire larval periods displaying nearly identical median lifespans to those treated with FUdR (whole larval period PX627_auxin = 17.7 days). However, auxin-exposed worms tended to display a lower rate of mortality late in life, yielding an overall significant difference between these treatments (CPH contrast: *z* = 3.04, *p* = 0.0089). The largest difference in lifespan was observed in PX627 individuals exposed to auxin during only the L4 stage of development, which had both longer median and maximum lifespans than matched FUdR treated individuals (L4 PX627_auxin = 19.2; CPH contrast: *z* = 5.76, *p* < 0.0001). Additionally, replicate trials from individuals treated only during the L4 stage tended to display more error variance (total variance attributable to replicate + plate effects) than the other experimental treatments (12% versus 2-4%, respectively).

**Figure 4.**
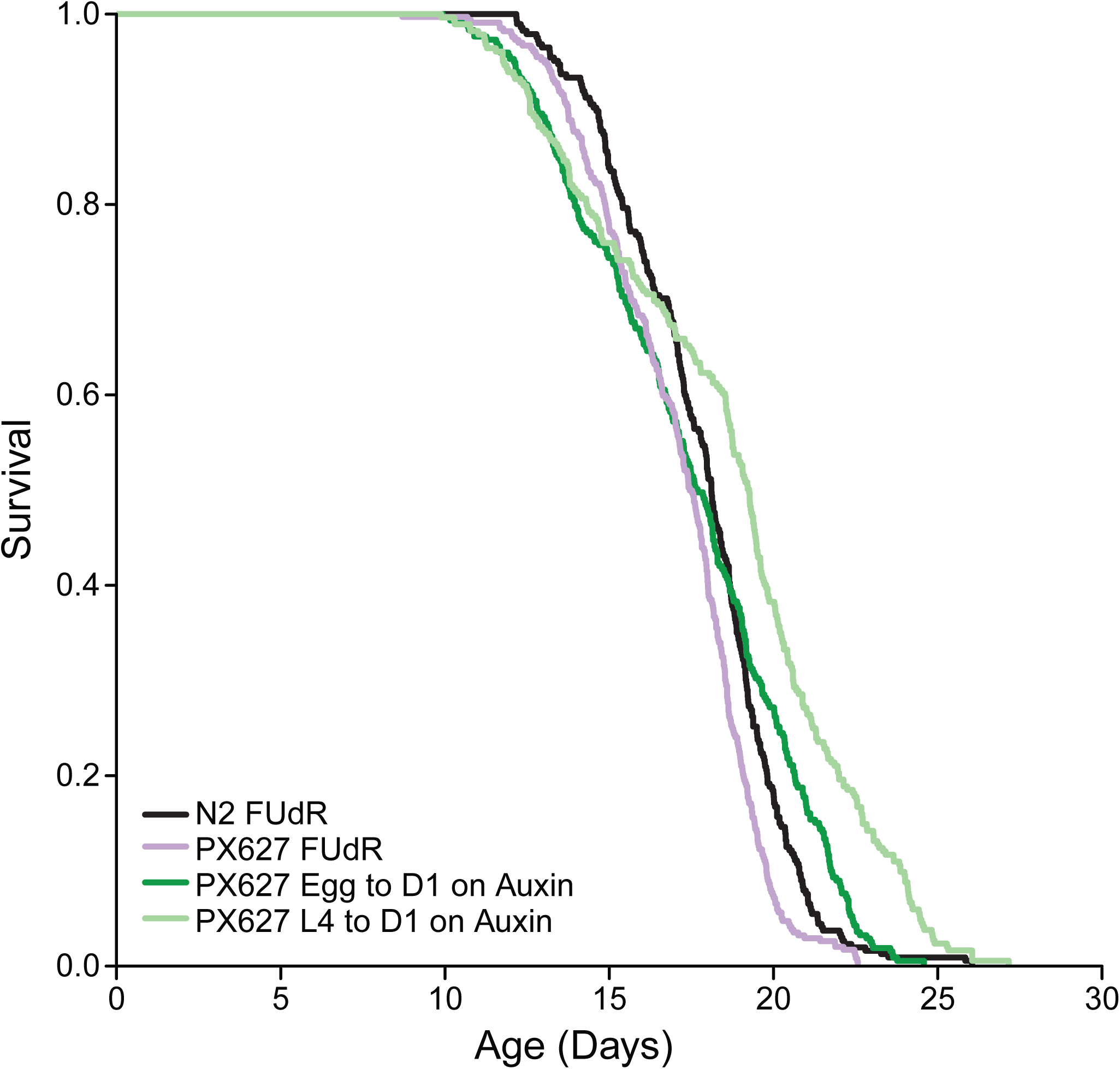
Lifespan curves comparing FUdR sterility to auxin-induced self-sterility. Wildtype (N2, black) and (PX627, purple) adults were FUdR treated. PX627 individuals were exposed to auxin from egg to day 1 of adulthood (dark green) or during the L4 stage alone (light green). Each survivorship curve represents six to eight pooled replicates each with over 100 individuals. The survivorship profiles were very similar across treatments and genetic backgrounds, though the PX627 L4 auxin treatment showed a quantitatively distinct profile.

## DISCUSSION

Sexual reproduction integrates multiple processes across an organism’s life, including the generation of gametes, the act of finding and securing mates, and the production of offspring. Each of these steps has an associated cost (Lehtonen *et al.* 2012). On top of these direct consequences, antagonistic interactions between the sexes during the process of mating as well as conflicts between parents and offspring can further exacerbate reproductive costs. Additionally, the interplay between reproduction and other major life history processes, such as aging and stress response, can add additional fitness trade-offs (Adler and Bonduriansky 2014). However, precise manipulation and quantification of reproductive trade-offs in an experimental setting has proved challenging.

### Sterility induction system

Using the AID system, we designed an external, non-toxic spermatogenesis arrest in *C. elegans*, resulting in hermaphrodite self-sterility and reversible male sterility. Hermaphrodite self-sterility could be induced through auxin exposure during the spermatogenesis developmental window alone. However, auxin exposure throughout larval development also induced complete self-sterility and had no noticeable effects on development (also see Zhang *et al.* 2015). Since this continued larval exposure required very little manual intervention, it is the preferred method for sterility induction, rather than multiple transfers of individuals on and off auxin during late larval development. Additionally, self-sterile hermaphrodites recovered their full fertility when mated with a wildtype male. In males, auxin exposure during larval spermatogenesis initiation alone was not sufficient to induce sterility, likely because transcription of *spe-44* is increasing going into L4 (Kulkarni *et al.* 2012). However, males could be fully sterilized using auxin exposure throughout larval development or early adult development. Moreover, males could recovery their fertility in an age-dependent manner, presumably again due to the transcription level of *spe-44*.

Our sterility induction system has a broad range of applications within the fields of spermatogenesis, sperm competition, and mating systems evolution. The temporal control over male sperm production allows for developments in our understanding of spermatogenesis, including the rate at which sperm are produced and the amount of sperm stored. Additionally, this temporal control could be co-opted for precise studies of sperm competitive behavior under multiple mating scenarios. A particularly interesting application of this system is the study mating systems evolution. For example, a genetically identical population could be simultaneously evolved under hermaphroditic and obligate male-female mating regimes. Alternatively, populations could be evolved to switch between mating regimes to better understand the genomic implications of these transitions.

### A new approach for aging studies

*C. elegans* is one of the premiere model systems for studying the biology of aging. The first life-extending mutations were discovered in *C. elegans* (Friedman and Johnson 1988; Kenyon *et al.* 1993) and since then this system has been used in hundreds of studies to investigate a wide variety of questions in aging research (reviewed in Park *et al.* 2017). In particular, a number of studies have shown that the reproductive state of an individual, especially those controlled by germline-soma signaling systems, can have important consequences for longevity (Shi and Murphy 2014; Angeles-Albores *et al.* 2017). From a practical standpoint, reproduction can greatly complicate longevity assays in nematodes. Since the age at first reproduction is much shorter than median lifespan, there is the potential for several generations to be living on a plate at the same time, even if one starts with an initial age synchronized cohort. In most longevity studies this problem is solved either by manually removing (“picking”) adults to fresh media every day, which is very labor intensive and prone to error, or using a chemical means to sterilize reproductive adults. The most common sterilization technique involves the use of FUdR, which disrupts DNA replication in proliferating tissues such as the germline. Actively poisoning a subject while trying to accurately track their health and lifespan is obviously less than ideal.

The sterility induction system developed here provides an ideal alternative to existing chemical sterilization approaches in *C. elegans* and other nematodes. First, worms only need to be exposed to auxin during their larval development and can then be transferred to regular media as adults. This exposure window cuts down on expense as well as the need to constantly replenish an environmental toxin throughout adult life. Further, there is no concern about potential interactions between the sterilization agent and other external treatments such as food quality or chemical interventions (Lucanic *et al.* 2017). Overall, we find that longevity trajectories of self-sterilized individuals are very similar to individuals raised on FUdR (Fig. 4). The only substantive difference that we observed was in individuals that had only been exposed to auxin during the L4 stage that immediately precedes sexual maturity. These individuals lived longer and displayed more variable outcomes than those that were exposed to auxin throughout their entire larval period. Potentially, even those these individuals are sterile, there may be some progression through spermatogenesis that has lifespan ramifications. This observation will require further study and provides an opportunity for deeper investigation of the relationship between reproduction and lifespan.

Overall, inducible sterility implemented during larval development followed by a transfer to standard media appears to be a viable, non-toxic, and more natural means of conducting long-term longevity studies with *C. elegans.* The only major disadvantage to this approach is the dependence upon the genetic background that we have constructed here or reconstructing the required degron system components in other genetic backgrounds. While these components may limit the applicability in some genetic studies, there is a very large advantage for direct environmental and/or chemical manipulation studies. Further, new aging related mutant screens might be initiated using PX627 as the parental strain.

### Conclusion

Our targeted approach of a critical spermatogenesis gene and the potential applications should in principle be transferable to other systems where auxin-induction is viable, such as *Drosophila* and zebrafish. Additionally, many other types of cell-specific arrests should be targetable using the auxin-inducible system. Now firmly within the era of CRISPR/Cas9 transgenics, targeted, external induction systems, such as the method presented here, are possible. When coupled with the power of automated assays and next-generation sequencing techniques, the field is poised to gain a wealth of information previously unattainable.

## Acknowledgements

We would like to thank C. Sedore for assistance with the automated lifespan set-up and data processing. We would like to thank S. Banse for advice and the Phillips Lab for constructive comments. This work was supported by the National Institutes of Health (training grant T32 GM007413 to KRK and R01GM102511 and R01AG049396 to PCP) and the ARCS Foundation Oregon Chapter (KRK).

